# The hidden treasure in your data: Phasing with unexpected weak anomalous scatterers from routine datasets

**DOI:** 10.1101/068403

**Authors:** Raghurama P. Hegde, Steven C. Almo, Udupi A. Ramagopal

## Abstract

Single wavelength anomalous dispersion (SAD) utilizing anomalous signal from native sulfurs, or other atoms with Z≤20, generally requires highly redundant data collected with relatively long wavelength X-rays. Here we present experiences with two proteins, where anomalous signal from serendipitously acquired surface bound calcium atoms with data redundancy as low as 10 was utilized to drive *de novo* structure determination. In both the cases, the calcium atoms were acquired from the crystallization solution, and data collection strategy was not optimized for exploiting the anomalous signal from these scatterers. The X-ray data were collected at 0.98Å in one case and at 1.74Å in the other (wavelength optimized for sulfur but anomalous signal from Ca was exploited for structure solution). Similarly, using a test case, we show that the data collected at ~1.0Å wavelength, where the f" value for sulfur is 0.28e, is sufficient for structure determination using intrinsic sulfur atoms from a strongly diffracting crystal. Interestingly, it was also observed that SHELXD was capable of generating a substructure solution from high-exposure data with a completeness of 70% for low-resolution reflections extending to 3.5Å, with relatively low redundancy (i.e., 6.0 fold). Considering the fact that many crystallization conditions contain anomalous scatterers such as Cl, Ca, Mn etc., it appears that the data from well diffracting crystals should be processed, with anomalous pairs kept separate, so as to detect any fortuitous signal that might be present, which could prove useful in either solving the structure *de novo* or in accurately assigning surface bound atoms.

## 1. Introduction

Direct determination of macromolecular structures from the X-ray diffraction data alone is a major goal for the crystallographic community. Seminal work by Hendrickson and colleagues demonstrated that anomalous scattering from native-sulfur atoms alone was sufficient to support *de novo* structure determination (Hendrickson, 1981). Interestingly, this daring sulfur-SAD experiment was performed long before the advent of density modification, and the maturation of detectors and software for maximizing signals from carefully formulated synchrotron data collection strategies. It took another ~20 years (Dauter *et al.*, 1999) for the crystallographic community to embrace the power of sulfur anomalous scattering for the *de novo* structure determination of proteins. Since then, several experiments have highlighted the necessity for highly redundant data to attain the accuracy required to use weak anomalous signal for phasing (e.g., (Dauter & Dauter, 1999, Dauter & Adamiak, 2001, Ramagopal *et al.*, 2003b, Debreczeni *et al.*, 2003, Uson *et al.*, 2003, Sarma & Karplus, 2006, Wagner *et al.*, 2006) and so on). A recent comprehensive study covering ~140 structures solved using sulfur anomalous signal further highlights the necessity of collecting accurate data (Rose *et al.*, 2015).

Single-wavelength anomalous dispersion (SAD) has become the method of choice for *de novo* determination of protein crystal structures (Hendrickson, 2014), with it accounting for 73% of the structures, determined by experimental phasing, deposited in the Protein Data Bank (PDB, (Berman *et al.*, 2000)) in 2013 (Bunkóczi *et al.*, 2015). The use of intrinsic sulfurs and Se-methionine derivatives for experimental phasing of protein structures has been extensively discussed (e.g., (Hendrickson, 1981, Hendrickson *et al.*, 1990, Doutch *et al.*, 2012, Kim *et al.*, 2013, Ramagopal *et al.*, 2003b, Sarma & Karplus, 2006, Dauter *et al.*, 1999). When it’s not possible to use the anomalous signals from intrinsic sulfurs, or Se-methionine derivatives cannot be prepared, derivatives of anomalous scatterers such as Zn, As, Mn, halide ions, halogenated fragments etc., have been exploited for the purpose of experimental phasing (e.g. (Dauter & Dauter, 2007, Kim *et al.*, 2013, Liu *et al.*, 2011, Ramagopal *et al.*, 2003a, Salgado *et al.*, 2004, Bauman *et al.*, 2016). Recently, interest in native-SAD experiments using longer wavelength X-rays has grown (Weinert *et al.*, 2015, Cianci *et al.*, 2016, Liebschner *et al.*, 2016), as it avoids the need to prepare Se-methionine or other derivatives. At wavelengths typically accessible at synchrotron beamlines and home sources, the anomalous signal from sulfur is weak. The use of longer wavelengths at a synchrotron source is expected to result in better anomalous signal, for example at 2.1 Å, f" for sulfur is 0.98e, close to double the value at 1.54 Å (0.55e) corresponding to a copper anode home source. However, most beamlines are not optimized to work at longer wavelengths, and the use of longer wavelengths limits data quality due to a number of factors, including the maximum resolution that can be achieved, increased absorption, radiation damage and possibility of harmonic contamination (Doutch *et al.*, 2012). The work presented here, although represents phasing from surface bound calcium atoms, the success of such phasing depends on the occupancy and orderliness of the surface bound atoms.

Although, longer wavelength is preferred for exploitation of weak anomalous signal from atoms below Z<20, initial diffraction experiment on a new protein always aims at getting the highest resolution data, even when the sequence identity of the protein with the known structures is below the “twilight zone” for molecular replacement. Generally, most beamlines are optimized to collect data around 1.0 Å, where the expected anomalous signal for anomalous scatterers such as calcium and sulfur is low. Many crystallization conditions contain metals such as Ca, Mn as well as chlorine ions, and it has been noted in a study on phasing of 23 proteins, that in 90% of the cases light-atom ions such as chloride, phosphate, sulfate, potassium, calcium were interacting with the protein molecule (Mueller-Dieckmann *et al.*, 2007). Anomalous signals from these serendipitous anomalous scatterers can potentially be used in experimental phasing of protein structures. However, the use of weak anomalous signals from adventitiously bound weak anomalous scatterers for phasing in macromolecular crystallography is seldom discussed. Herein we discuss two such cases where anomalous signal from surface-bound calcium atoms acquired from mother-liquor were exploited for phasing: (i) PSPTO_5518 from *Pseudomonas syringae* pv. Tomato (**PROT1**) and (ii) hypothetical protein PTO0218 from *Picrophilus torridus* (**PROT2**). We also examined hen egg white lysozyme (**PROT3**), where data collected at ~1.0 Å wavelength was sufficient to drive *de novo* structure solution using sulfur anomalous signal alone. To the best of our knowledge, this is the first case where sulfur anomalous signal at ~1.0 Å was shown to be sufficient for *de novo* structure determination. Also, for PROT1, the data were collected near the selenium edge and for PROT2 near the iron edge; moreover, the redundancy was near or below 10 in all the cases. However the anomalous signals from these atoms, with Bijvoet ratio being at or below the Wang’s limit of 0.6% (Wang, 1985) in two cases (PROT1 and PROT3) and around 1% in the case of PROT2, could nevertheless be used to obtain the structures. These results suggest that a careful analysis of supposedly native data and a little manual help can lead to the unanticipated determination of structures.

## 2. Materials and Methods

PROT1 was crystallized using hanging drop vapor diffusion, from a drop containing 7.8 mg/ml protein solution mixed in 1:1 ratio with reservoir solution consisting of 30% PEG 400, 0.1 M HEPES, 0.2 M calcium chloride, pH 7.5, maintained at 293.0K. PROT2 was crystallized using sitting drop vapor diffusion, from a drop containing 10 mg/ml protein solution mixed in 1:1 ratio with reservoir solution consisting of 0.2 M calcium chloride dihydrate, 0.1 M HEPES - Na pH 7.5, 28% v/v polyethylene glycol 400 maintained at 298K. PROT3 was purchased from Sigma-Aldrich and used without further purification. It was crystallized using hanging drop vapor diffusion from a drop containing 20 mg/ml protein solution mixed in 1:1 ratio with reservoir solution consisting of 10% sodium chloride, 50mM sodium acetate, pH 4.6, maintained at 293K.

Diffraction data for PROT1 and PROT2 were collected at the X4A beamline at NSLS using a ADSC Quantum 4 CCD detector. Diffraction data for PROT3 were collected at X29A beamline at NSLS (Brookhaven National Laboratory) using an ADSC Quantum 315 CCD detector. The data were processed using HKL2000 (PROT2) (Otwinowski & Minor, 1997) and HKL3000 (PROT1, PROT3) (Minor *et al.*, 2006). All data were collected from crystals maintained at 100K in a stream of cold nitrogen gas. CC_1/2_(anom) for each of the datasets was calculated as follows: each dataset was split into two equal sets and this was treated as MAD data, with one set input as peak data and the other set input as inflection data in HKL2MAP to calculate the anomalous correlation coefficients and these were used as CC_1/2_(anom).

For all the three proteins, the SHELX program suite (Sheldrick, 2010), as incorporated in HKL2MAP GUI (Pape & Schneider, 2004), was used for experimental phasing. Substructure solution, followed by phase calculation, was performed using SHELXC/SHELXD, with 1000 cycles of substructure search, E value cut-off of 1.7, unless otherwise mentioned, to include only strong reflections, and density modification in SHELXE. The density modified maps were input to ARP/wARP (Langer *et al.*, 2008) for model building. For PROT2, the quality of electron density map obtained after density modification alone was not sufficient for model building and three cycles of iterative chain tracing with 20 cycles of density modification was used. To check the feasibility of an automated structure solution workflow, the anomalous data were input to the AutoSol routine in PHENIX (Adams *et al.*, 2010).

PROT1 and PROT2 did not have model in PDB that were appropriate for molecular replacement, and the datasets described here were used for phasing, as well as refining the structures. Coordinates have been deposited in the PDB as 2PAG and 2I52 for PROT1 and PROT2 respectively. For lysozyme (PROT3), the coordinates from 1LZ8 were used as a reference structure. Since in all three cases, we had nearly complete models from phasing, the phases that yielded the model were refined against the final deposited structure (moving the final structure to the same origin as the experimental model), with 20 cycles of restrained refinement in REFMAC5 (Murshudov *et al.*, 2011), to obtain the final phases. The initial and final phases were merged together into one file using the program CAD(CCP4), and the average phase errors before and after density modification (DM) were calculated using the program PHISTATS (CCP4) using final refined map as reference. Similarly, map correlations before and after density modification were calculated using the program overlapmap (Braenden & Alwyn Jones, 1990, Jones & Stuart, 1991). All these programs are available as part of the CCP4 software suite (Winn *et al.*, 2011).

## 3. Results

Table 1 presents the diffraction data for each of the three proteins and Table 2 the absorption edges, with corresponding energies, of the anomalous scatterers encountered in this study. In the following subsections the phasing approach used for each of these proteins is described.

**Table 1.**
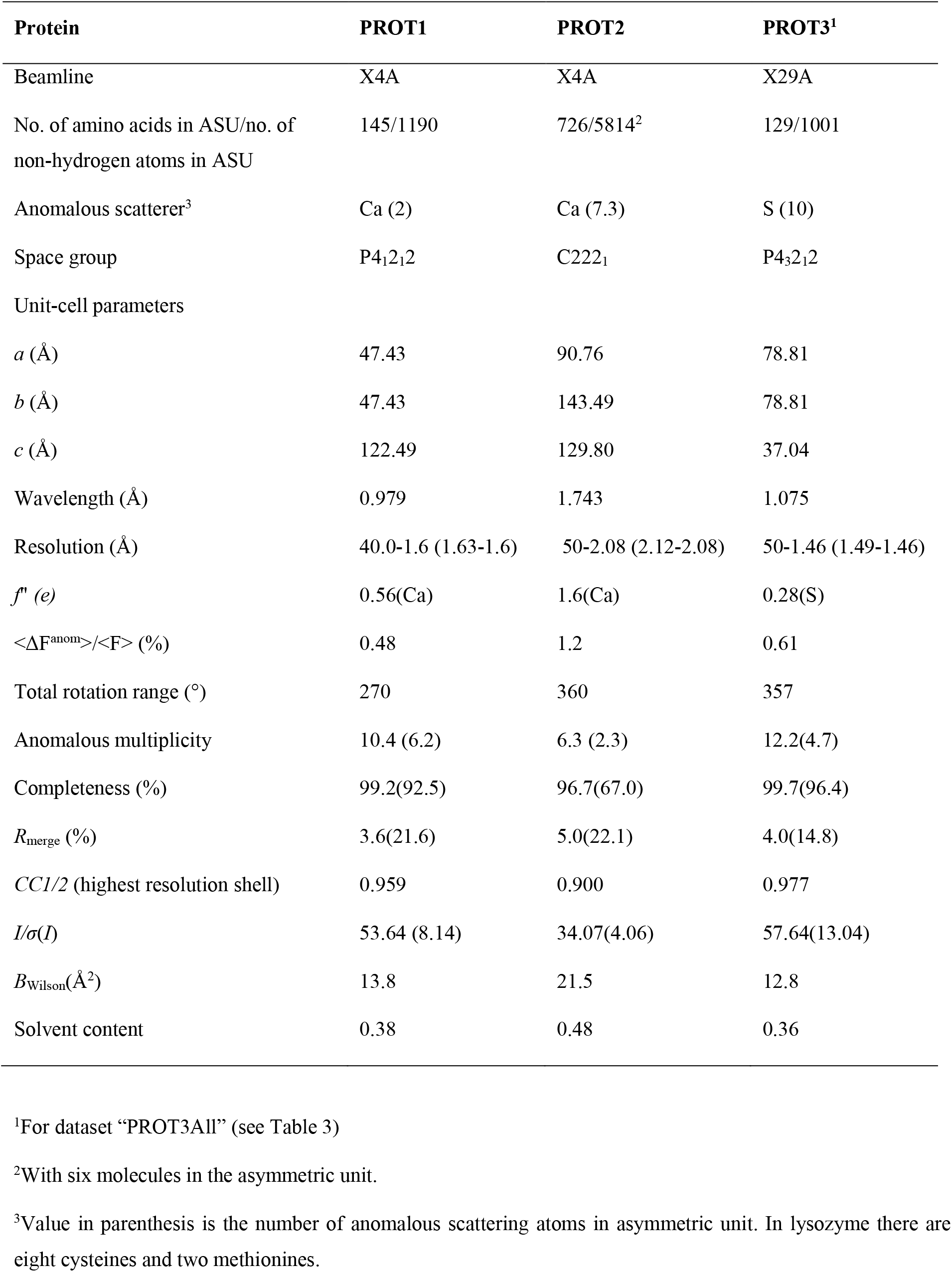
Diffraction Data. (Unless otherwise mentioned values in parenthesis are for the highest resolution shell)

**Table 2.** Absorption edges of anomalous scatterers observed in this study and the difference between absorption edge energy and energy at the wavelength used for data collection

| Absorption edges |  |  | Difference from absorption edge and the data collection wavelength (in keV) |  |  |
| --- | --- | --- | --- | --- | --- |
| Atom | Wavelength (Å) | Energy (keV) | Protein | Energy used (keV) | Difference from absorption edge(keV) |
| Ca | 3.070 | 4.038 | **PROT1** | 12.664 | 8.626(Ca) |
| S | 5.016 | 2.472 | **PROT2** | 7.113 | 4.043(Ca) |
|  |  |  | **PROT3** | 11.533 | 9.061(S) |

### 3.1. PRO1

PROT1 crystallized in the space group P4_1_2_1_2 with one molecule in the asymmetric unit and diffracted to 1.6 Å. This protein has 3 cysteine residues. BLAST search (Altschul *et al.*, 1990) against the PDB did not reveal any model, suggesting the sequence is highly unique and ruled out the possibility of determining the structure by Molecular Replacement. Native X-ray diffraction data were collected at the wavelength 0.979Å with a crystal rotation of 0.5° per frame and a total of 540 frames were collected covering 270° wedge, with an exposure time of 10 seconds. Although, the intention was to collect accurate and high-resolution native dataset, it was our practice to check for presence of anomalous scatterers in most high-resolution datasets (>1.8 Å). It should be noted that scaling the data in HKL2000 with the “scale anomalous” option did not indicate the presence of any anomalous signal (see sub-section 4.2 and Figure 3). This observation is not surprising as the sulfur edge is almost ~10,200 eV from the energy of the X-rays used in this experiment. However, to our surprise, a substructure search with SHELXD looking for four anomalous scatterers, using anomalous data up to 2.1 Å, produced two strong peaks followed by four additional peaks, which are consistent in most correct solutions (Figure S1). These substructure coordinates, along with phases from SHELXD, were input to SHELXE for density modification and phase improvement. Phases obtained from SHELXE were input to ARP/wARP for model building, resulting in a model containing residues 4-135 of the 145 amino acid protein. The two strong peaks in the substructure solution suggested an anomalous scatterer slightly larger than sulfur. The crystallization condition contained calcium chloride, from which calcium atoms could have been acquired. On refinement of the model two strong difference density peaks were observed on the surface near Gln(101), Asp(103) and Thr(105). The coordination with the amino acid side chains and water molecules together with anomalous difference Fourier map suggested these were probably bound calciums (Figure 1(a)), which were modeled in the refined structure.

**Figure 1.**
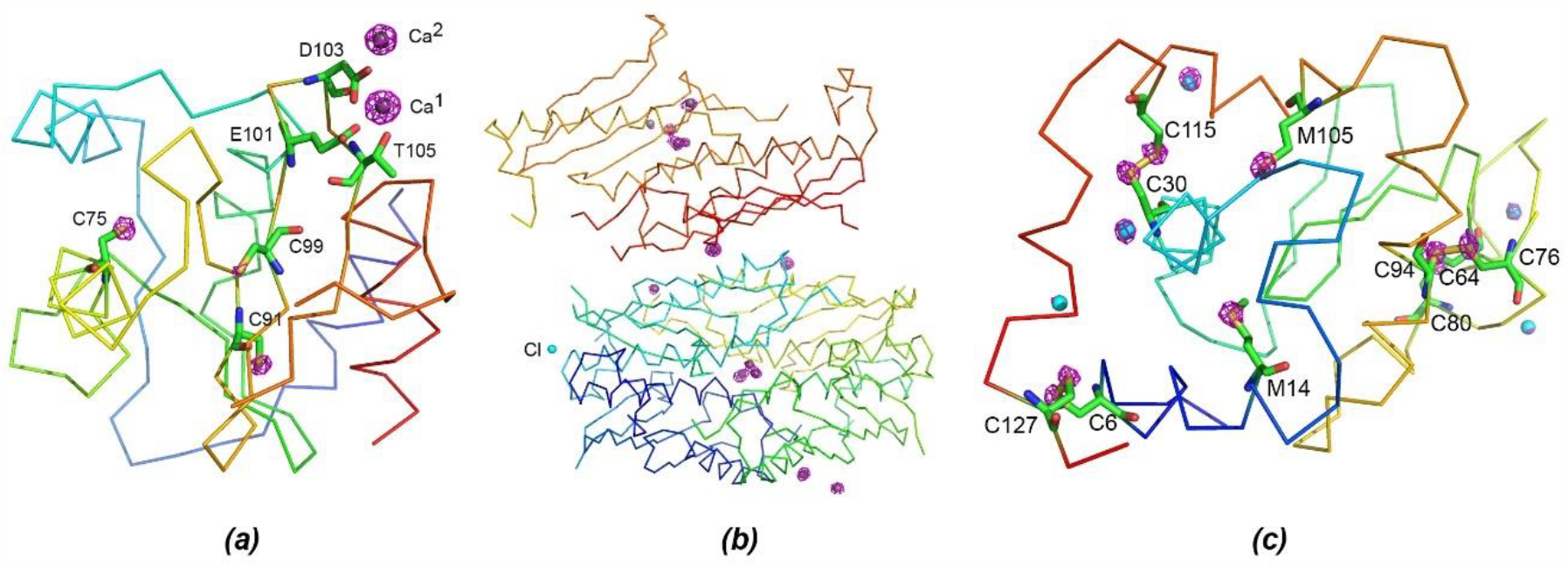
Representation of the structures of the three proteins with anomalous maps displayed around the anomalous scatterers in the proteins. The protein chains are shown as C^α^ traces and the anomalous maps are shown as purple meshes contoured at 5σ level. (a) PROT1, the two calcium atoms are shown as grey spheres, the residues coordinating with the calcium atoms and the cysteines in the protein chain are shown and labelled; (b) PROT2, the calcium atoms are shown as grey spheres and the chlorine atom as a cyan sphere. Each of the six chains in the asymmetric unit are colored differently; (c) PROT3, the cysteines and methionines in the protein chain are shown and labelled, the chlorine atoms are shown as cyan spheres. Made using PyMOL (Schrodinger, 2010).

To examine the efficacy of a completely automated crystal structure determination workflow, the anomalous data were input to the AutoSol routine in PHENIX. Interestingly, when the anomalous scatterers were input as two calcium atoms, PHENIX was able to build from residues 4 to 111 and 124 to 136; but, when the anomalous scatterers were input as 4 sulfurs it was able to build only 53 residues - 7-24, 72-80, 86-106, 126-130, about 37% of the structure. The modeled segments correspond to correct elements of the structure; but, PHENIX appeared unable to build a more complete model. However, using either calcium or sulfur as input to SHELXD yielded the same substructure solution and in both the cases the density-modified map was good enough for complete automatic building.

### 3.2. PROT2

PROT2 is a 13kDa hypothetical protein PTO0218 from *Picrophilus torridus.* Again, a BLAST search did not identify a useful molecular replacement model. PROT2 crystallized in the space group C222_1_ with six molecules in the asymmetric unit and diffracted to 2.08Å. PROT2 contains 121 residues, including one cysteine and four methionines. X-ray diffraction data were collected at 1.743Å (7,113 eV), near the iron edge with a crystal rotation of 1° per frame, with the goal of determining the structure by sulfur phasing. As the data were collected at a bending-magnet beam line X4A, each frame required four seconds of exposure. Given the speed of SHELXD (and thanks to HKL2MAP), there was sufficient time to check the substructure solution at frequent intervals. After approximately 300 frames, substructure solution looking for 30 sulfurs yielded several solutions with high CC/PATFOM in SHELXD (Figure S1). A significant drop in occupancy was observed between the first and second heavy atom peaks, and between the second and third, with a smooth transition in occupancies of additional sites. In general, for substructures involving Se or S atoms (i.e., covalently bonded to the protein), such variation in occupancy is rare. The observed drop in the occupancy between the third and fourth atom suggested the presence of anomalous scatterers other than sulfur. The initial phases obtained from SHELXD using best substructure solution and anomalous data up to 2.8 Å, were input to SHELXE for 20 cycles of density modification with tracing option switched-off. The maps obtained with the original and inverted substructures exhibited a clear contrast in terms of connectivity, CC and FOM. These phases were input to ARP/wARP and automatic model building failed. To improve the initial phases, SHELXE was rerun with 3 iterative cycles of chain tracing subsequent to 20 cycles of density modification, which enabled ARP/wARP to build 678 of the 726 residues.

The asymmetric unit in the crystal of PROT2 consists of four molecules arranged in a rugby ball-like shape and the other two adopting a half rugby ball-like shape positioned parallel to the long side of the ball (Figure 1(b)). To exploit non-crystallographic symmetry (NCS) information in density modification, NCS-operators were used in the program DM (Cowtan & Zhang, 1999), and the resulting map was input to ARP/wARP for model building, allowing 659 residues in 12 chains to be built (the structure 2I52 was originally determined exploiting NCS). To examine the efficacy of a completely automated crystal structure determination workflow, the anomalous data were input to the AutoSol routine in PHENIX, which was able to build 640 of the 726 residues.

In the final structure there were 12 calcium atoms, which refined with varying occupancies. Two calcium atoms were refined with full occupancy and the rest with partial occupancies, with a total occupancy of calcium atoms of 7.3 (which is around one Ca per chain). All the calcium atoms, which were derived from the crystallization media, are in the solvent region, either in between the molecules or on the surface. One of two calcium atoms that refined with full occupancy mediated a crystal contact.

Although this case represents a dataset optimized for phasing using sulfur anomalous signal, anomalous signal from surface bound calcium atoms contributed significantly to phasing and *de novo* structure determination of this protein. PROT1 represents a dataset where anomalous signal from surface bound calcium atoms at 0.979 Å were utilized for phasing whereas PROT2 represents a dataset where anomalous signal from surface bound calcium atoms at 1.743Å is utilized. Hence, despite not being a typical routine dataset, we included PROT2 dataset in this study along with PROT1 to compare the quality of the phases obtained in the two data sets. (discussed further in section 4.2).

### 3.3. PROT3

The two above cases prompted us to test the feasibility of structure determination using sulfur anomalous signal from a “routine” dataset collected without optimization for sulfur phasing. The beamline X29 at NSLS employs a mini-gap undulator source, with maximum flux observed around 11.5 keV (wavelength ~ 1.078Å, (Shi *et al.*, 2006)). As a test case, we used the Hen egg white lysozyme (HEWL), a well-studied 129 amino acid protein with 8 cysteines and 2 methionines (referred to as PROT3). At this wavelength, the f" value for sulfur is 0.28 electrons and the expected anomalous signal is as low as 0.61%. The protein crystallized in the space group P4_3_2_1_2 with one molecule in the asymmetric unit and diffracted to 1.46 Å. X-ray diffraction data were collected at 1.075Å (11.533 keV) with a crystal rotation of 1° per frame and 0.5 second exposure. Owing to the strong diffraction, 200 frames were collected with the detector at ~200mm (corresponding to ~ 1.46Å) with a 7-fold attenuation, followed by another 157 frames with unattenuated beam, both with a crystal rotation of 1°. These two datasets were merged and used for phasing. Substructure solution was attempted searching for 14 sulfur atoms to include surface bound chlorine atoms (observed in most structures with crystallization condition containing NaCl), with three disulfide bonds resolved (DSUL 3, command in SHELXD), using anomalous data up to 2.0 Å. Considering the fact that (1) the data were collected approximately 9000 eV away from the sulfur edge, (2) the total wedge for the both the datasets together is only around 360 and (3) the f" value at this wavelength for sulfur is very low (0.28e), we did not anticipate successful identification of the sulfur positions. However, out of 1000 cycles, 3 solutions with high CC/PATFOM and clear bimodal distribution distinguishing the correct and wrong solutions (Figure S1) gave a clear indication that the substructure solution was successful. Coordinates of the first few anomalous scatterers in all three putative solutions were manually checked for consistency of the sulfur positions and the coordinates from the best solution were used for phasing in SHELXD followed by density modification with SHELXE with solvent content of 38%. As expected for lysozyme, the map obtained from the original coordinates (space group P4_3_2_1_2) after 20 cycles of density modification had better CC, contrast, connectivity and FOM, as compared to the one obtained from the inverted coordinates substructure coordinates (space group P4_1_2_1_2). The resulting map was input to ARP/wARP for model building; a single chain of 126 residues was docked in to the map.

To determine the minimal data needed for phasing, different datasets were prepared using (the dataset obtained with unattenuated beam is referred to as high intensity data and the one obtained with attenuating the beam is called low intensity data): (1) first 100 frames from the high intensity data merged with last 100 frames from the low intensity data, (2) first 150 frames from both datasets merged together, (3) first 100 frames from both datasets merged together, (4) all 200 frames of the low intensity data and (5) all 157 frames of high intensity data alone. Table 3 provides a comparison of the various parameters for these datasets. Henceforth PROT3All refers to the dataset obtained from merging all of the high and low intensity datasets. We were able to successfully phase and build a model with all the datasets. For datasets (3) and (4), 20 cycles of density modification alone was not able to produce a map that was good enough for ARP/wARP to build a model, but 3 iterative cycles of tracing with 20 cycles of density modification yielded a map that ARP/wARP could use to build 126 of 129 residues of the protein chain. Datasets (1) and (3) have similar multiplicity and resolution, but the total wedge covered in dataset (1) is 200°, whereas it is 100° in dataset (3). Similarly, the higher quality of the initial phases from dataset (1) compared to dataset (4) highlights the importance of merging the high-intensity data, which is expected to increase overall accuracy of the data.

**Table 3.**
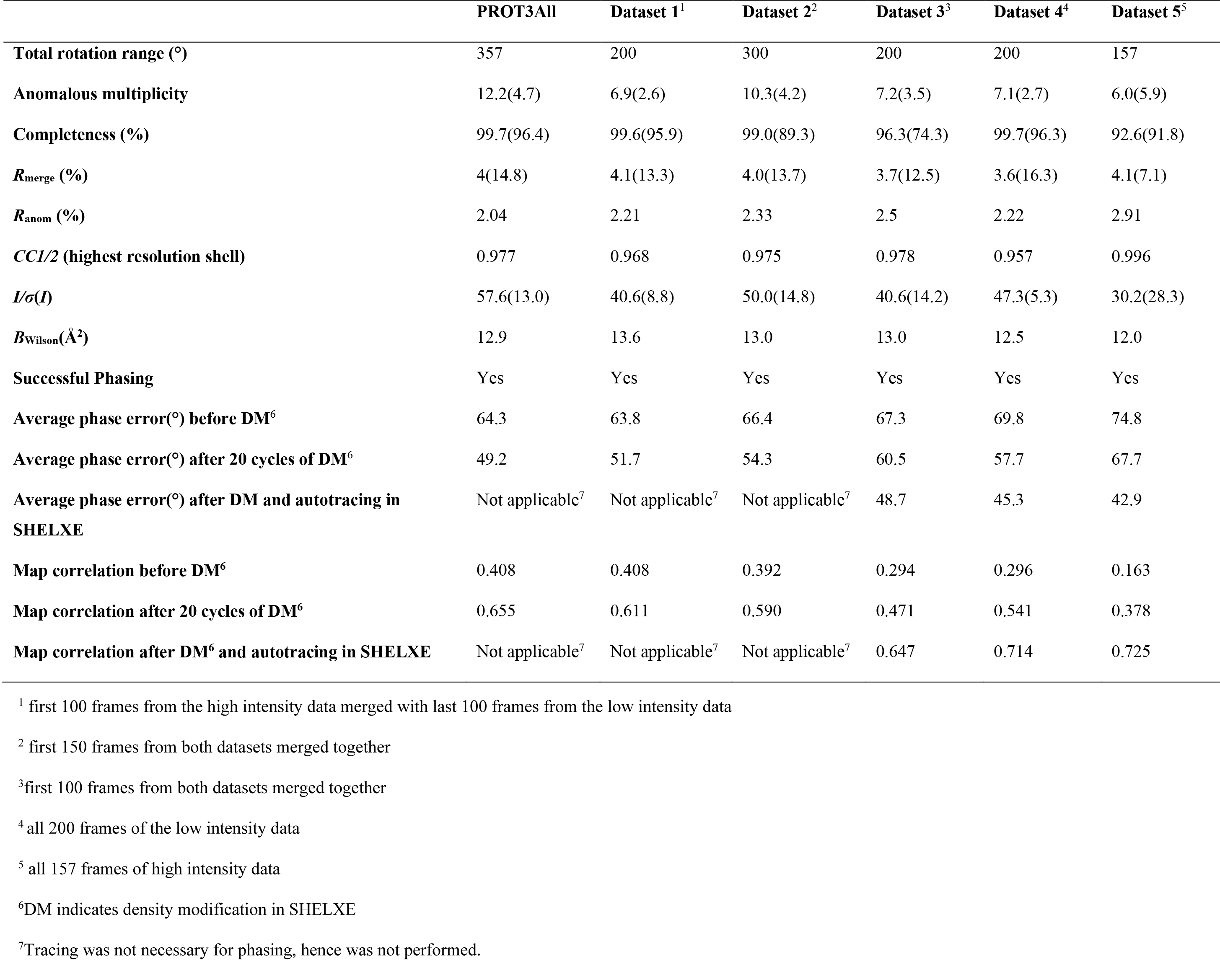
Comparison of PROT3 datasets. (values in parenthesis are for the highest resolution shell)

Determining the substructure solution using dataset (5) required several attempts that involved tuning the E values (normalized structure factor) and resolution. In one of such attempts, using anomalous data extending to 1.7 Å with an E value cut-off of 1.7 gave one correct solution out of 1000 cycles (Figure 2). The difficulty in finding the substructure solution is not surprising considering the fact that this data had completeness of only ~70% in the range 50-3.5 Å (Figure S2) due to a high number of overloads and redundancy as low as 6.0 (Table 3). These missing high-intensity reflections are expected to carry a disproportionate amount of information about the structure.

**Figure 2.**
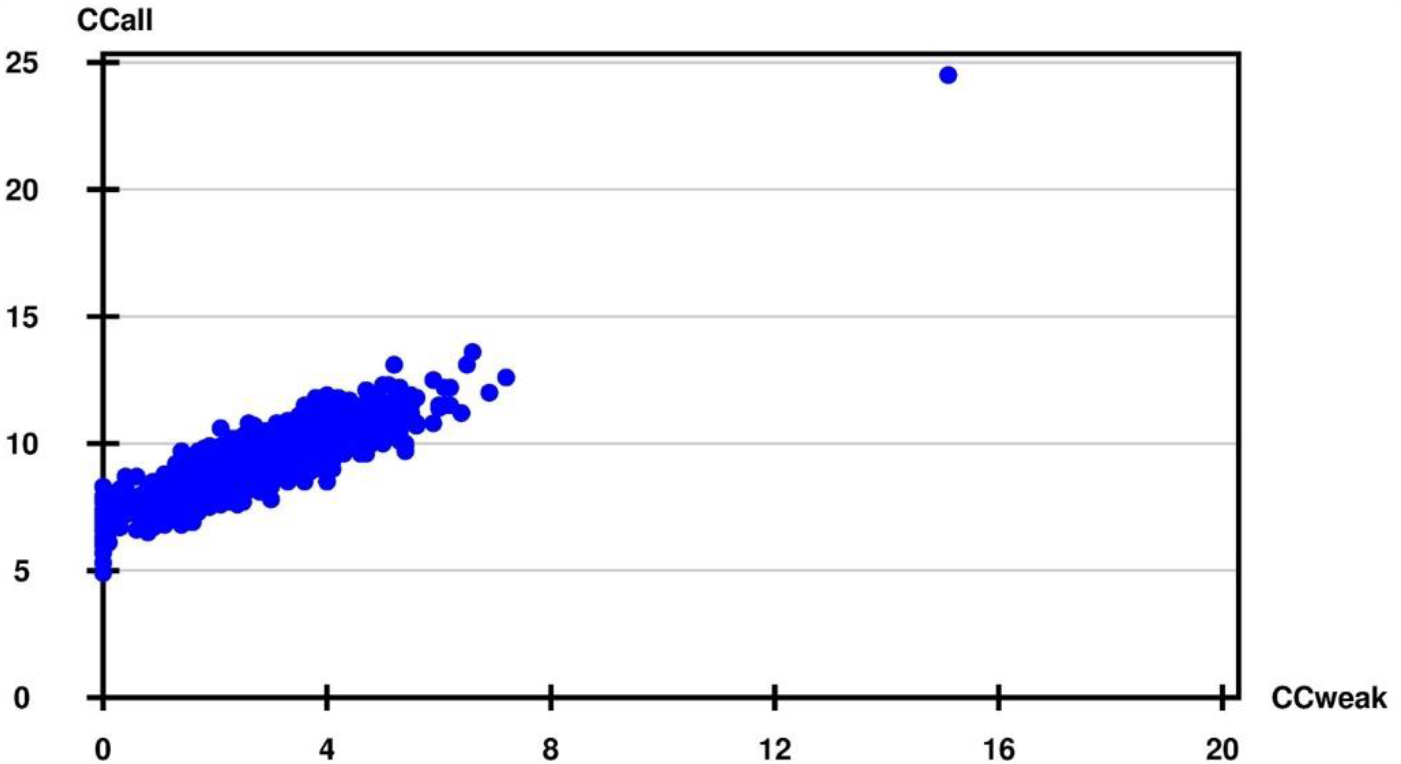
CCall/CCweak from SHELXD run of 1000 cycles representing 999 wrong substructure solutions and one correct solution that resulted in successful phasing of Dataset 5 for PROT3, which has a data completeness of only ~70% till 3.5Å due to overloads, with average redundancy as low as 6.0.

Interestingly, Autosol in PHENIX was not able to produce a correct substructure solution with the complete dataset PROT3All and hence we were not able to solve the structure with this program using default run. Based on our previous experiences, we believe with correct substructure PHENIX should be able to produce similar quality map.

## 4. Discussion

As mentioned earlier, PROT1 and PROT2 had no models in PDB and *de novo* structure determination was necessary in both the cases. The data acquisition strategies were not optimized for anomalous scattering, as these crystals unexpectedly acquired anomalous scatterers from the crystallization solution. In addition, our experiment with test case PROT3 was performed to mimic the most routine experiments that are used to acquire native datasets. In the cases of PROT2 and PROT3 (high intensity data alone, see Table 3), we were able to determine structures with redundancies as low as 6.0; for PROT1 this value is around 10.0. Moreover, the actual energy of the X-rays used for data collection were ~8,500 eV (PROT1), 4000 eV(PROT2) and 9000 eV(PROT3) away from the resonance edges associated with the relevant anomalous scatterers in PROT1, PROT2, PROT3 (Table 2). All three cases mimic data collection strategies typically employed for the collection of native data. Despite not being optimized for experimental phasing these datasets yielded very good quality phases sufficient for automatic model building.

### 4.1. Redundancy, radiation damage and data accuracy

It is accepted fact that high redundancy is one of the key factors for successful sulfur phasing, even when X-ray energies corresponding to higher f" values (e.g., 5-8 keV) are used. One must be mindful that redundancy comes with the detrimental effects of radiation damage-induced errors (Garman & Nave, 2002, Holton, 2009, Ravelli & Garman, 2006, Garman, 2010). A dose limit of 2 × 10^7^ Gy (Gy= J.kg^−1^) was proposed by Henderson for cyro-cooled protein crystals, at which the intensities of diffracted rays are reduced by half (D_50_) (Henderson, 1990). Based on a recent experiment, although this limit is relaxed to 3 × 10^7^ Gy or 30 MGy where the average diffraction intensities reduced to 70% of original value (D_70_) (Owen *et al.*, 2006). Another recent study (Liebschner *et al.*, 2015) suggests that the damage rate may be different for different crystals and indicates that the damage D_70_ was 30% more at 6.33 keV (7.5 MGy) as compared to 12.66 keV (11 MGy), highlighting that longer-wavelengths result in greater damage. The acceptable extent of damage depends on the objective of the experiment, based on which, quality of the data obtained requires a different level of accuracy; for example, whether the experimenter is looking for overall structure or critical biological information where highly radiation-sensitive residues like aspartate or glutamate are involved or exploitation of anomalous information to solve the structure *de novo* and so on. A series of 20 consecutive datasets collected from a single crystal of Thaumatin, covering the same rotational range of 0-90° at a second-generation bending magnet beam line, showed intensity variations as great as 300% for some reflections (Banumathi *et al.*, 2004). In this case, overall dose received by the crystal was approximately 3.4% of Henderson limit for each dataset (covering only 90° wedge). These data indicate that even a relatively small received dose can cause as much as a 10-15% variation in the intensity of some reflections between consecutive datasets, although not all reflections may be affected to this extent. Allowed variation due to radiation damage is much more stringent for the case of anomalous phasing with very weak anomalous signal. Liu et al., proposed that for multi-crystal native-SAD, a dose of 5MGy is the upper limit for a dataset from each individual crystal(Liu *et al.*, 2014). In the case of PROT1, PROT2 and PROT3 the overall dose, as calculated by the program *RADDOSE* (Zeldin *et al.*, 2013), are 0.10, 0.03 and 0.13 MGy, which is 0.5 %, 0.15% and 0.65% of the Henderson limit, respectively. For the low-intensity dataset of PROT3, this value is only 0.02 MGy. These values indicate that the crystals experienced very minimal damage, and consequently the accumulated intensity error due to radiation damage is also minimal. It should be noted that while collecting data for multi-crystal native SAD, the variation due to non-isomorphism may not be insignificant; however, it has been shown that by properly choosing and merging compatible datasets, structures can be determined from weakly diffracting crystals (Liu *et al.*, 2014). It appears that the accumulated error resulting from radiation damage is more serious than slight non-isomorphism between crystals.

Errors due to X-ray beam line instabilities are also detrimental to data quality. Multiple measurements of the same and symmetry-related reflections can improve overall accuracy of the data. Accuracy of the measurement also increases with the strength of the diffraction data. As shown in Table 1, the overall I/σ(I) values are as high as 50 for the PROT1 and PROT3 datasets. In these cases, although the expected anomalous signal and the redundancies are low, accurate measurement of very strong reflections and their Friedel mates, as well as overall low X-ray dose, appears to compensate for the low signal and low redundancy.

As described above, two sweeps of data were collected for PROT3, one with seven-fold attenuation covering a 200° wedge and the other unattenuated covering 157° wedge. As expected, substructure solutions from merged data had higher overall CC_all_/CC_Weak_ and PATFOM as compared to other low redundancy datasets. Although, there were a few overloads in the low intensity data, overall, the data completeness was close to 100% (Figure S2). To determine the minimum data required for determining the PROT3 (HEWL) structure and to examine whether the high intensity data contributed to successful phasing, we merged various wedges from the low and high intensity data. The redundancies of these datasets were around 7.0 in most of the datasets (Table 3) and phasing almost universally successful, except for the dataset where only high-intensity data was used, where almost 30% of data in the low resolution till 3.5 Å were missing due to overloads; overall I/σ(I) for this data was as low as ~30, whereas it is 50 for the low exposure data. Although, substructure solution was difficult when using the high intensity dataset alone (dataset 5), the quality of the map was comparable to other PROT3 datasets (Table 3). These observations highlight contribution of missing reflections to overall I/σ(I) which in turn could determine the ease/success of phasing.

### 4.2. Anomalous signal and phasing

Several indicators for the estimation of anomalous signal have been proposed (Zwart, 2005, Dauter, 2006). While such indicators are very useful in most phasing experiments, they may not be fully accurate when dealing with very weak anomalous signal. One such indicator, χ^2^-statistics provided by data reduction program HKL2000, is very useful for detecting the presence of anomalous signal at the time of data collection itself. However, for very weak anomalous scatterers with Z≤20, χ^2^-statistics may not fully report on the extent of anomalous signal available (Figure 3). In a multi-crystal native SAD experiment, it was observed that data with higher multiplicity from multiple crystals produces better substructure solutions and final maps, as compared to datasets with better values for some of these indicators, including CC_1/2_, but lower multiplicity (Liu *et al.*, 2014).

**Figure 3.**
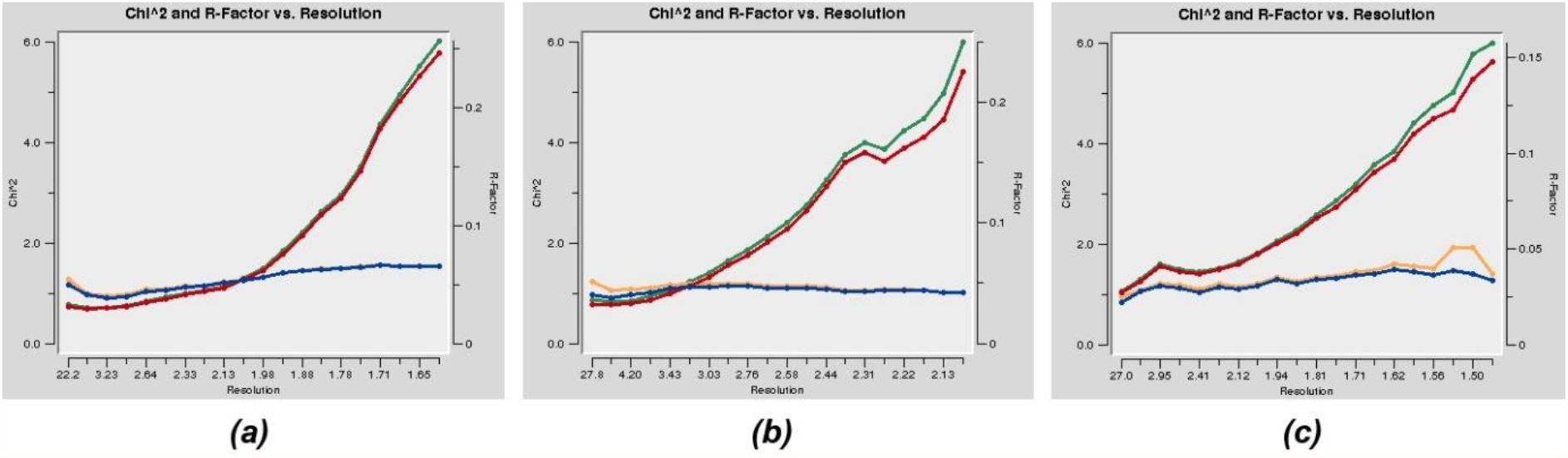
χ^2^ vs resolution plots for (a) PROT1, (b) PROT2 and (c) PROT3. The blue and red lines represent χ^2^and R-factor respectively for the data with Friedel mates separated, orange and green lines represent χ^2^ and R-factor respectively for the data with Friedel mates merged.

In all the three cases discussed above, none of these tests were performed before determining the structures. Analysis of these datasets suggests that detectable anomalous signal was present in all cases. Figure 4 shows the <d"/σ> values as calculated by SHELXD plotted against resolution for all the proteins along with the values corresponding to SeMet derivatized putative glucarate dehydratase from *Acidaminococcus sp. D21*(PDB id 4HYR), as representative of data collected for an anomalous scatterer at an energy closer to its peak (Se in this case). The CC_1/2_(anom) values for all the three datasets clearly indicate significant anomalous signal, albeit modest, extending to at least 3.0 Å resolution (Figure 5). Since the datasets were collected at 1.74 Å and 0.979 Å respectively for PROT2 and PROT1 (f" values of 1.6 and 0.56 respectively for Ca atoms), it was expected that higher CC_1/2_(anom) values would be observed for PROT2 as compared to PROT1. Surprisingly CC_1/2_(anom) did not vary between the two datasets (Figure 5), underscoring the suggestion that in cases with very weak anomalous signal, this metric might not fully predict the likelihood of succeeding in substructure solution and phasing. However, consistent with CC_1/2_(anom) values both the datasets produced map of similar quality (Table 4).

**Figure 4.**
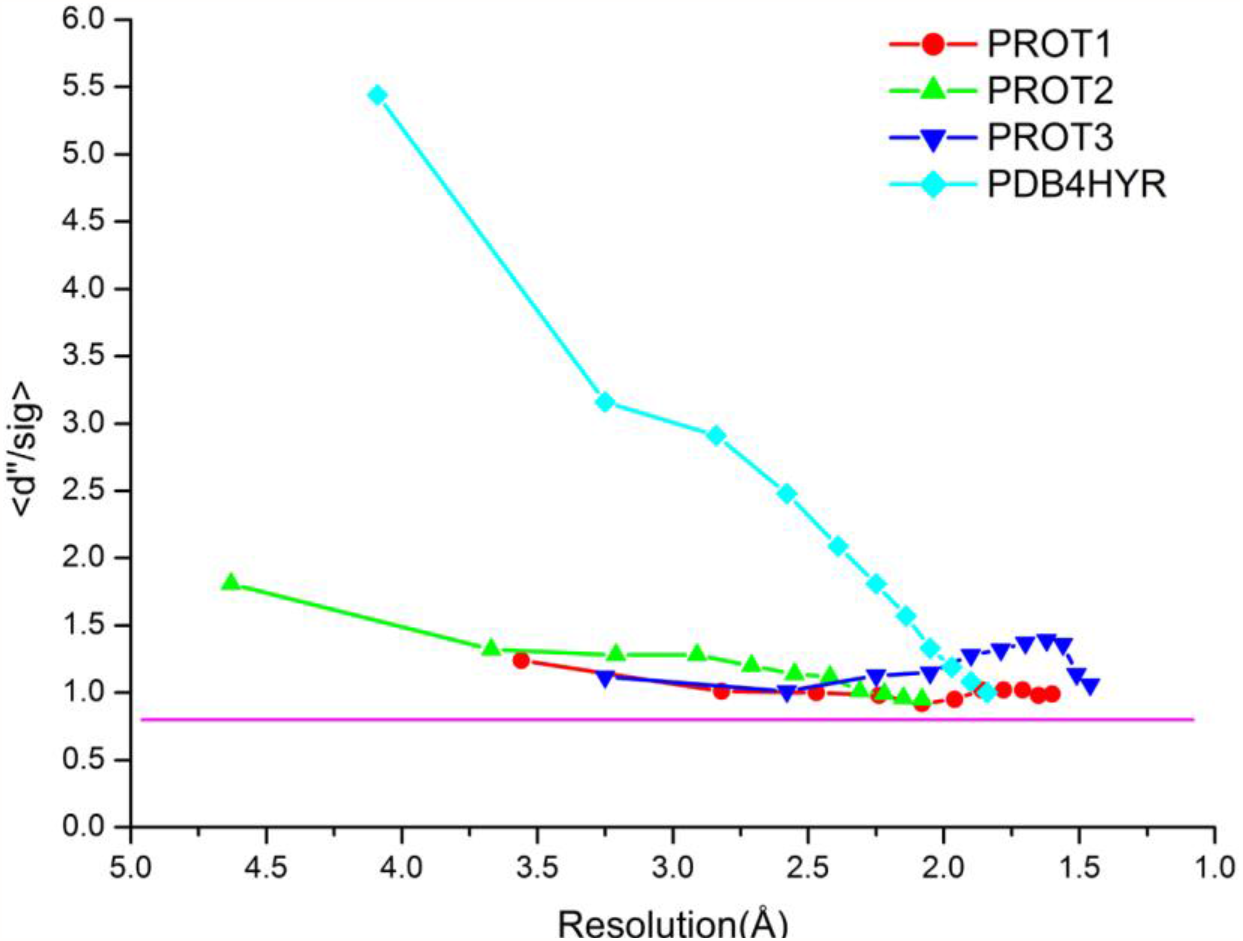
A plot of <d"/sig> vs resolution for the three proteins discussed in this paper along with a representative selenomethionine data collected at Se edge of a putative glucarate dehydratase from *Acidaminococcus* sp. D21, PDB id. 4HYR. The values for PROT3 correspond to the dataset PROT3ALL. The purple line is drawn at <d"/sig> = 0.8 about which the signal is considered to be zero. All the three proteins show weak anomalous signal throughout the resolution range.

**Figure 5.**
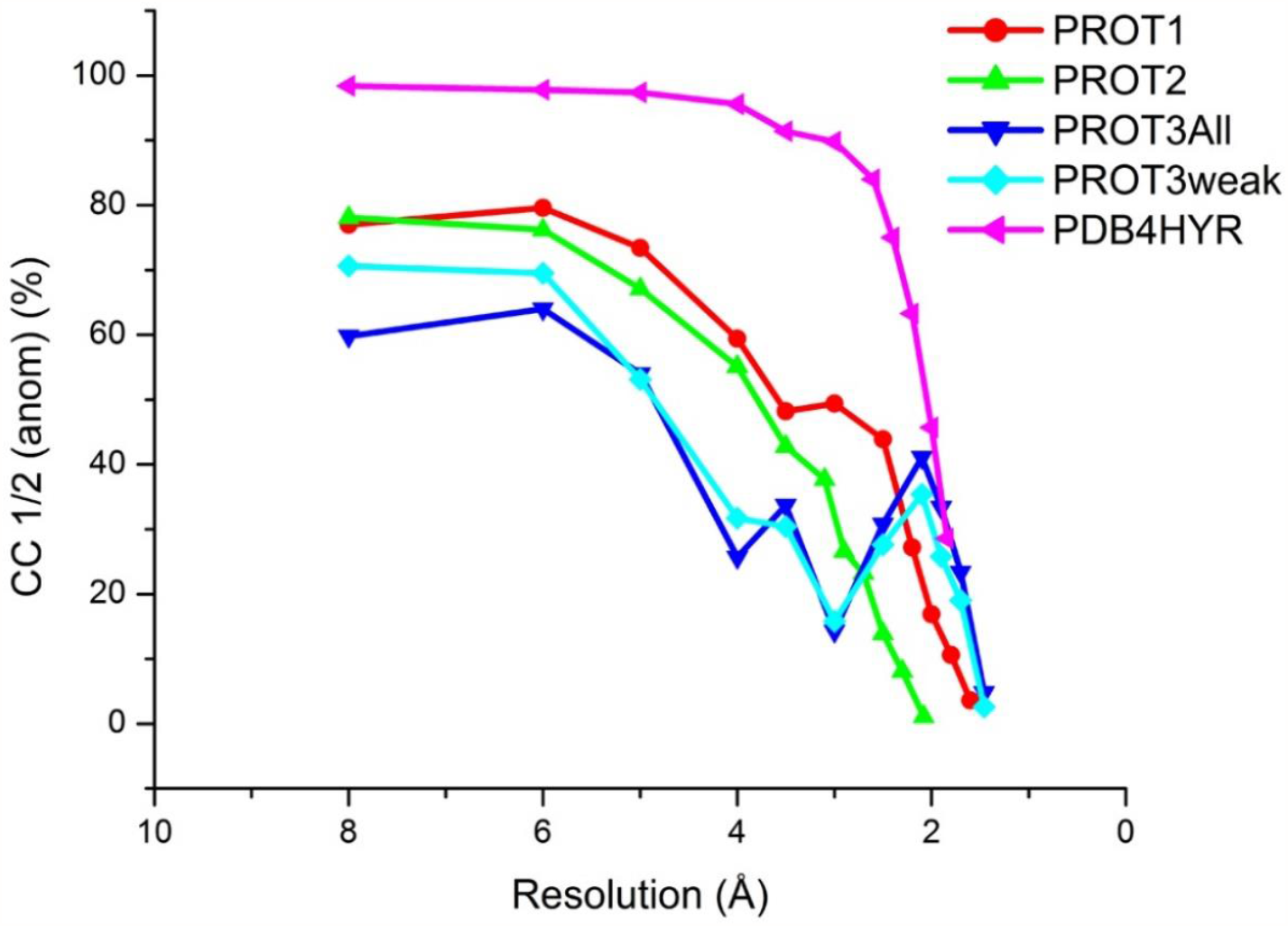
A plot of CC_1/2_(anom) vs resolution for the three proteins discussed in this paper with a representative selenomethionine data collected at Se edge of a putative glucarate dehydratase from *Acidaminococcus* sp. D21, PDB id. 4HYR. For PROT3 CC_1/2_ (anom) vs resolution is plotted for both the dataset with high intensity and low intensity data merged (PROT3All) and for the weak intensity data alone (PROT3weak, corresponds to Dataset 4 in Table 3).

**Table 4.** Map correlation coefficients and phase errors

| Protein | Map correlation |  | Average phase error (°) |  | Solvent content |
| --- | --- | --- | --- | --- | --- |
|  | before DM<sup>1</sup> | after DM<sup>2</sup> | before DM<sup>1</sup> | after DM<sup>2</sup> |  |
| PROT1 | 0.382 | 0.539 | 71.4 | 61.1 | 0.38 |
| PROT2 | 0.391 | 0.596 (0.730)<sup>3</sup> | 70.6 | 61.5 (49.8)<sup>3</sup> | 0.48 |
| PROT3<sup>4</sup> | 0.396 | 0.6 | 66.8 | 52.7 | 0.36 |

<sup>1</sup>Comparison between initial phases before density modification in SHELXE and the final phases (from the refined model)
<sup>2</sup>Comparison between phases after density modification in SHELXE and the final phases.
<sup>3</sup>Autotracing in SHELXE was required before these could be phased successfully. Value in parenthesis is the phase error/map correlation after 3 cycles of autotracing with 20 cycles of density modification before each cycle of autotracing in SHELXE.
<sup>4</sup>Corresponds to PROT3All dataset.

Similarly, it is notable that the data for PROT2 was collected at a wavelength of ~1.743 Å and the substructure contained ~7 calcium atoms (12 atoms with 10 of them refined with partial occupancy), which has an f" value of 1.6e at this wavelength. The quality of the phases obtained from this data is not superior to those obtained from PROT3All dataset (Table 4), where the wavelength for data collection was 1.075 Å and anomalous contribution is from sulfur atoms, which has an f" value of 0.28e at this wavelength. Both these datasets have similar redundancy, although the calculated <ΔF^anom^>/<F> (%) value (Table 1) for PROT2 is twice of that for PROT3. However, PROT3 has higher overall I/σ(I) (Table 1) indicating that while collecting redundant data is critical for successful phasing, redundancy should not be achieved at the expense of overall data strength. It should be noted that most successful cases of multi-data native-SAD experiments at low-resolutions were from strongly diffracting crystals having overall I/σ(I) of around 50 and having high symmetry (Weinert *et al.*, 2015, Liu *et al.*, 2014).

The observed strength of the anomalous signal from a given scatterer not only depends on the wavelength of the X-ray and the number of scatterers but also on the B-factor of the anomalous scatterers and quality of data. The impact of B-factor can be clearly seen in the case of PROT3 dataset. Although, the expected f" values for chlorine and sulfur atoms are 0.36e and 0.28e, respectively, the first surface bound chlorine atom was observed as a seventh peak in the substructure solution of PROT3All data (Table S1), the other five chlorine atoms were not consistent in all the solutions; they either appeared after all the sulfur atoms in the substructure solution or did not appear at all. Anomalous difference Fourier peaks for PROT3 dataset is shown in Figure 6 at 3, 5 and 7σ levels. It is clear that some of these surface bound chlorine peaks can’t be seen in the map contoured at 5σ level (cyan spheres in Figure 6(b)), while the anomalous peaks of the sulfurs, which are part of the protein chain and hence more ordered than the chlorines, are observed.

**Figure 6.**
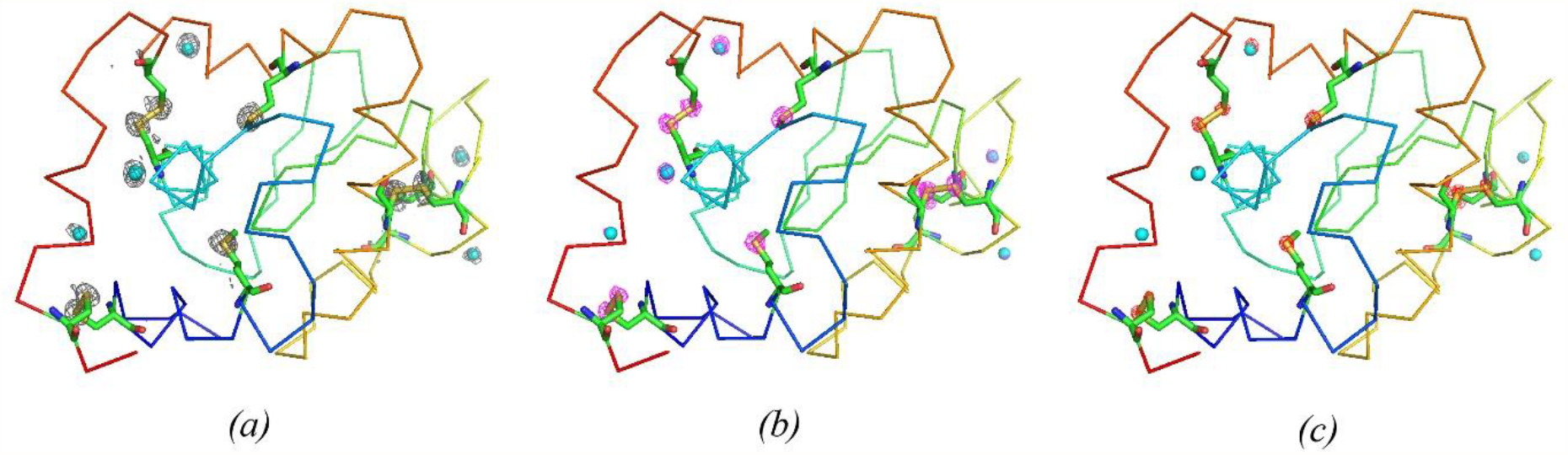
Anomalous maps around the sulfurs and chlorine in PROT3 shown at (a) 3σ, (b) 5σ and (c)7 σ. The maps are calculated using the PROT3ALL dataset. The sulfurs in the side chains are in yellow and the chlorine atoms in cyan. Anomalous difference Fourier map at 7σ is not seen around most chlorine atoms. Made using PyMOL (Schrodinger, 2010).

## 5. Conclusions

The exploitation of very weak anomalous signal is becoming an increasingly common practice and native SAD with redundant data collected from multiple crystals or multiple datasets from the same crystal but at different locations of the crystal has been shown to work with crystals diffracting to lower than 3.0 Å (El Omari *et al.*, 2014, Liu *et al.*, 2014, Akey *et al.*, 2014). In most cases it is essential to have highly redundant data (with 20-100 multiplicity) collected using long-wavelength (1.5 to 2.5Å) X-rays to maximize the accuracy and strength of the anomalous signal. Here, we show that routinely collected high-resolution datasets from strongly diffracting crystals, using X-rays of shorter wavelength (around 1.0 Å) with f" values of 0.28e for sulfur atoms and 0.56e for Ca atoms can drive the *de novo* structure determination. These can be considered as routine datasets in the sense that the wavelengths used were not optimized to maximize the anomalous signal, neither was the data collected with very high multiplicity for SAD phasing.

Whether it is for anomalous phasing or for molecular replacement, processing data with anomalous pairs kept separate has at least two advantages (1) calculation of anomalous difference Fourier map with refined phases would help in the accurate assignment of bound atoms. In two of the cases presented here, although the wavelength used is around 1.0Å, anomalous difference Fourier peaks varied between 30-5σ (Table S1) depending on the anomalous scatterer and the B-factor of these atoms. It appears that strongly bound atoms with Z>15 can be assigned from the strength of the anomalous difference Fourier peaks together with coordination geometry/environment around the bound atoms, even from data collected using ~1.0Å wavelength. (2) In fortuitous cases, unexpected anomalous signal can be used for phasing the structure *de novo* where the starting model is of poor quality. In all the three cases presented here, the phase error before density modification were as high as ~70 to 75° which is close to random phase assignment of 90°, suggesting the initial phases information required is no longer limiting and the current density modification and improvement protocols can start from the phases that approach random phases. Hence, with much more accurate detectors of today, effective phasing programs, highly efficient *ab initio* programs like ARCIMBOLDO (Rodriguez *et al.*, 2009, Rodriguez *et al.*, 2012) and with the combination of phase information from various sources (maybe for e.g., NCS to pull the phase error away from 90°), it is tempting to propose that we are moving closer to determining most structures from strongly diffracting native crystals without intended derivatization.

## Acknowledgements

Ramagopal likes to thank the Department of Biotechnology, Government of India for the award of a Ramalingaswami fellowship and the Vision Group on Science and Technology, Government of Karnataka, India for an infrastructure grant.

